# *Drosophila melanogaster* as a model host to study arbovirus–vector interaction

**DOI:** 10.1101/2020.09.03.282350

**Authors:** Mandar S. Paingankar, Mangesh D. Gokhale, Deepti D. Deobagkar, Dileep N. Deobagkar

## Abstract

Arboviruses cause the most devastating diseases in humans and animals worldwide. Several hundred arbovirus are transmitted by mosquitoes, sand flies or ticks and are responsible for more than million deaths annually. Development of a model system is essential to extrapolate the molecular events occurring during infection in the human and mosquito host. Virus overlay protein binding assay (VOPBA) combined with MALDI TOF/TOF MS revealed that Dengue-2 virus (DENV-2) exploits similar protein molecules in *Drosophila melanogaster* and *Aedes aegypti* for its infection. Furthermore, the virus susceptibility studies revealed that DENV-2 could propagate in *D. melanogaster*, and DENV-2 produced in fruit fly is equally infectious to *D. melanogaster* and *Ae. aegypti*. Additionally, real time PCR analysis revealed that RNAi coupled with JAK-STAT and Toll pathway constitutes an effector mechanism to control the DENV-2 infection in flies. These observations point out that *D. melanogaster* harbors all necessary machineries to support the growth of arboviruses. With the availability of well-established techniques for genetic and developmental manipulations, *D. melanogaster*, offers itself as the potential model system for the study of arbovirus-vector interactions.

## INTRODUCTION

The molecular and biochemical basis of the invertebrate host specificity to arboviruses is interesting but comparatively less understood phenomenon. Most of the virus/parasite interacting proteins are house keeping in nature^1–5^ (also see Supplementary Information Table S1). For example, actin, Heat shock proteins, prohibitin and tubulin are reported as arbovirus interacting proteins in *Aedes* cells^2,3,5–7^. These polypeptides are present ubiquitously in other insects also. As parasites/viruses are dependent on their symbionts for reproduction, we might expect that it would be advantageous for parasites/viruses to have broad host ranges and an abundance of potential vectors. Susceptibility, early multiplication and production of virulent *Plasmodium gallinaceum* in non-vector non hematophagous insect *Drosophila melanogaster* support this hypothesis^8^. Arboviruses infect more than 100 million people worldwide every year. The cellular mechanism for transmission and the complex molecular interplay between arboviruses and their vectors are not well characterized. It has hampered development of novel strategies for disease intervention and control. Genomic sequences are available for various diseases causing vectors in literature and Genomic databases^8^. Molecular mechanisms associated with virus propagation in mosquito have been studied using genomics approach in *Ae. aegypti* and *Anopheles* mosquitoes^10–12^. In these two species, the availability of the whole genome sequences helped in identification of possible molecular mechanisms involved in virus infections. However, in many other insect vectors such as ticks, sand flies, fleas and other members of Culicidae family, limited specific proteomics information limit the in depth understanding of vector parasite interaction in these species. Considering the sequence and function similarities in the receptor polypeptide components, house keeping proteins, innate immune system genes and other regulatory gene sequences in the eukaryotes, it could be feasible to establish a test system for the study invertebrate parasite/virus interactions.

The *Drosophila* model system has been explored to study a variety of human infections and diseases^12–18^. Furthermore, the utility of *D. melanogaster* as a host model system for understanding cellular interactions of various human pathogens has been well-established^8,19–28^. However, *D. melanogaster* underutilized in arbovirus studies and holds great potential in the understanding mechanisms involved in arbovirus pathogenesis. Therefore, as a step towards understanding the molecular mechanisms involved in arbovirus-vector interactions, the present work seeks to develop a *Drosophila* model of dengue infection that better reflects the molecular events in the human and mosquito infections.

In the current study, we report that *D. melanogaster* can serve as a useful model system for the growth and the propagation of DENV-2. The virus produced in *D. melanogaster* can also infect the *Ae. aegypti* similar to the virus grown in mosquito. We report that the infectivity and multiplication of the virus grown in *D. melanogaster* and *Ae. aegypti* is comparable. *Drosophila* model system has biosafety advantage over *Ae.aegypti*, as *Drosophila* do not feed on blood and never transmit any infectious diseases though bite. Therefore, the application of this model system thus could also be extended to the other arboviral infection analysis.

## MATERIAL AND METHODS

### Ethics statement

Rules laid by Institutional Animal Ethics Committee (IAEC) affiliated with National Institute of Virology (NIV), Pune, India were followed for handling of animals. These experiments were carried out in a biosafety level-2 facility of the NIV. All animal experiments were approved by the IAEC and experiments were performed as per the guidelines laid by the Committee for the Purpose of Control and Supervision of Experiment on Animals (CPCSEA), India.

### Drosophila flies and <u>Ae</u>. <u>aegypti</u> mosquito

Oregon K stocks of *D. melanogaster* were grown on standard cornmeal–agar medium at 24°C. *Ae. aegypti* mosquitoes were reared in laboratory conditions at 28±1 °C, 70±5% relative humidity (RH) and light: dark (LD) 12:12 h. Adult mosquitoes and flies, 3–6 days of age were used in the infection experiments.

### Virus stock preparation

Virus stock was prepared by inoculating TR1751 strain of DENV-2 in mice by following the procedure described earlier^29^. Virus titer of randomly picked vial was determined by plaque assay (8.23× 10^6^ PFU/ml).

### Plaque assay

The DENV-2 titer in the flies and mosquitoes was calculated using plaque assay method as described earlier^29^. The virus titer in the carcasses individual mosquito (virus blood fed or virus injected) or fly (virus injected) was reported as plaque forming units (PFU) (values are expressed as the mean ± SD).

### DENV-2 infection

Infections were carried out by injecting ~1 μl of a viral suspension (8.23× 10^6^ PFU /ml) into the thoraces of *D. melanogaster* adult flies (n=246) and *Ae. aegypti* (n=168). Actual injected titer of DENV-2 was determined using plaque assay at 2h p.i.. Infected flies were then maintained at 24° C. *Ae. aegypti* mosquitoes were infected with DENV-2 via blood through membrane feeder (n=76) and then maintained at 24°C for 11 days. To check infectious nature of DENV-2 produced in *D. melanogaster*, carcasses of DENV-2 positive flies were crushed and centrifuged at 4°C, 10000 rpm, for 30 min and supernatant filtered through 0.22μm syringe filter and inoculated in *D. melanogaster* (n=94). The homogenates were mixed with blood and oral fed to *Ae. aegypti* (n=82). *Antibodies*

DENV-2 was inoculated in three-four weeks old Swiss albino mice intra-peritoneally and booster doses of DENV-2 were given (one dose/week) along with Freund’s incomplete adjuvant (1:1) for two weeks. The mice were injected with 10% ascitic tumor cells intra-peritonealy. The intra-peritoneal fluid collected and after removal of the debris by centrifugation, the supernatant was used to check for the presence of antibodies. Anti DENV-2 antibodies were incubated with the mosquito midgut extract to remove the non-specific antibodies. The protein-A column was used to purify the antibodies. The pre-immune serum was collected and checked for presence of non specific antibodies against the DENV-2.

Monoclonal antibodies against *D. melanogaster* anti-beta actin antibody [mAbcam 8224] - loading control [ab8224] (Abcam, USA), anti-tubulin antibody [T0950] (Sigma, USA), Monoclonal Anti-Hsc70 (Hsp73) antibody [SAB3701436] (Sigma, USA), anti-HSP70 antibody ([5A5] (ab2787), Abcam, USA) and anti-prohibitin antibody [EP2803Y] (ab75766) (Abcam, USA) were used to confirm the results obtained in the various assays.

### Detection of dengue viral antigen in the head squashes

The presence of viral antigen was determined by indirect immunofluroscence assay (IFA) in the head squashes of mosquitoes/flies as described by Apte-Deshpande *et al*.^29^. Along with each experiment, positive and negative controls were processed using the same protocol. Presence of DENV-2 antigen detected in head squash preparation of *D. melanogaster* flies on everyday till 10 days post infection (p.i.).

### Membrane Fraction isolation

Brush-border membrane fractions (BBMF) from guts of *Drosophila* were isolated according to the procedure described earlier^2^.

### Virus Overlay Protein Binding Assay (VOPBA)

VOPBA was performed to identify cell polypeptides involved in virus binding following the procedure described earlier^2^. Experiments were performed independently four times and negative controls (without virus incubation, without antibody incubation) were kept. In VOPBA, interacting proteins were identified, therefore no positive control is available for this assay.

### Protein identification using MALDI-TOF/TOF MS

Bands corresponding to DENV-2-binding activity were excised from gels and were subjected to reduction, alkylation, followed by in-gel digestion with trypsin. Extracted peptides were desalted using the column and were separated on a Biobasic C18 capillary column. The chromatographically seperated peptide masses were analyzed by matrix-assisted laser desorption/ionization time of flight (MALDI-TOF/TOF) on Ultraflex TOF/TOF (Bruker Daltonics, Germany). The proteins were identified using the mass spectrum produced from each sample by searching the m/z values against the protein databases (NCBInr, MSDB, and Swissprot) using the MASCOT, MSfit and Profound search engine. Parameters used for identification of proteins were fragment ion mass tolerance of 0.40 Da, parent ion tolerance of 0.4 Da and iodoacetamide derivative of cysteine as a fixed modification. The monoclonal anti-prohibitin antibody ([EP2803Y, ab75766, Abcam, USA), anti-tubulin antibody (T0950 Sigma, Aldrich, Germany), anti-HSP70 antibody ([5A5] (ab2787), Abcam, USA) were used to validate the results obtained in the MALDI TOF/TOF MS analysis.

### RNA isolation

The QIAamp viral RNA mini kit (Qiagen USA) was used to isolate the viral RNA from carcasses of flies. Total RNA was isolated from DENV-2 positive carcasses of *Ae. Aegypti* and *D. melanogaster* flies using the RNA purification kit (Ambion-Thermo Fisher USA).

### Detection of DENV by RT-PCR

Detection of DENV-2 in *D. melanogaster* and *Ae. aegypti* was performed using a RT-PCR procedures described earlier^30^. The viral RNA was converted into cDNA using Goscript cDNA synthesis system and the PCR amplification was performed in a Veriti thermocycler (Life technology, USA). Negative controls consisted of RNA from uninfected *Ae. aegypti* and *D. melanogaster* flies and water instead RNA. The second round of PCR was performed with 2 μl of sample from first round of amplification reaction.

### Real-time qPCR assays

RNA samples (2μg) were incubated with Turbo DNase (Ambion, USA) and reverse-transcribed using High capacity cDNA synthesis system (Life technologies, USA). Real-time relative quantification of 50 ng of cDNA was carried out using the Power SYBR Green PCR Kit (Life technologies, USA) and ABI Detection System ABI 7300 (Applied Biosystems, USA). Four independent biological replicates were conducted for each sample which were loaded in duplicates. Primer sequences for Drosophila genes were retrieved from Flyprimerbank^31^ and are listed in Table 1. Fluorescence detection was performed at the end of each extension step and amplicon specificity was checked by dissociation curve analysis at a rate of 1°C every 30 s from 60 to 95°C. All samples were amplified in duplicate from the same RNA sample and the mean value was calculated and was used for relative fold change analysis. The quantitative expression of the target gene was normalized to 18s mRNA in the same samples.

**Table 1:**
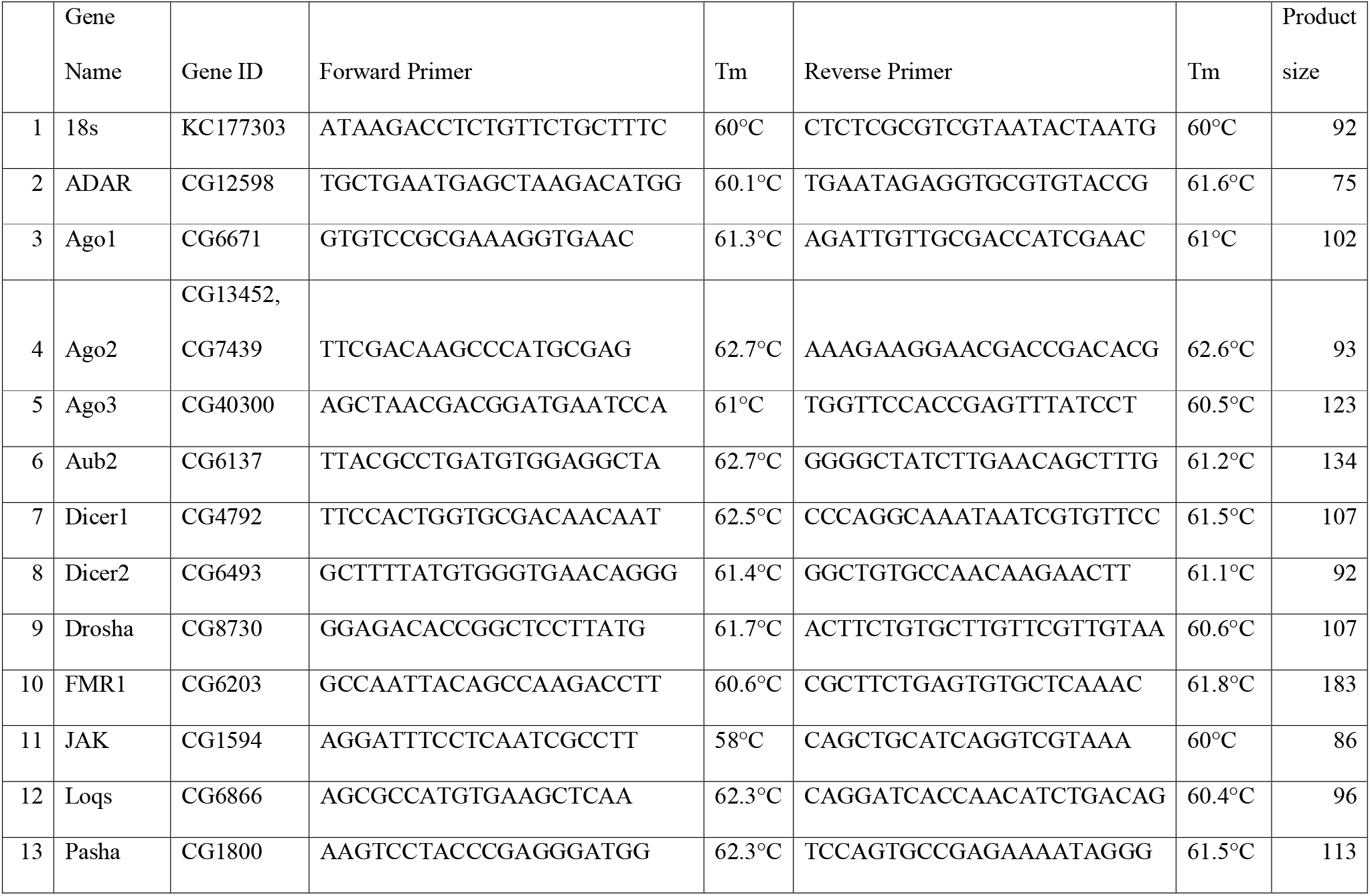
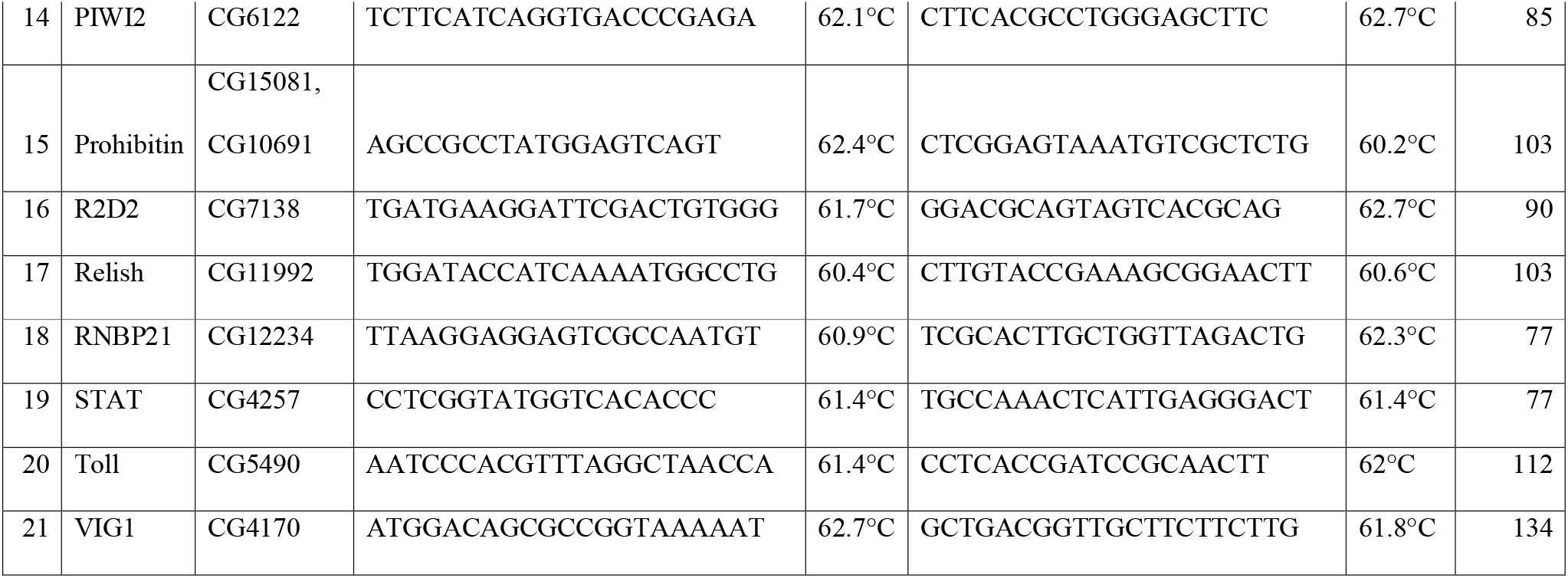
Primers used for real time PCR.

### 2.15 Statistical analysis

The groups (control vs infected, *Aedes aegypti* DENV-2 injected vs *Drosophila* DENV-2 injected, 2 hrs p.i. vs 7 days p.i., fold change in control vs infected flies) were compared by nonparametric Mann-Whitney U test. The viral loads were log-transformed and were compared by non-parametric Mann-Whitney U test.

## RESULTS

### Does DENV-2 exploit similar molecules in different insects?

The studies on DENV interacting proteins in *Aedes* cells suggest that housekeeping molecules are exploited by DENV to establish the infection. Therefore, we identified the *D. melanogaster* gut BBMF proteins which are interacting with DENV-2 using one dimensional and two dimensional VOPBA and MALDI TOF/TOF analysis.

When immobilized brush border membrane fraction polypeptides were incubated with DENV-2, seven polypeptides, Belle, gamma-tubulin ring complex subunit, HSP 70Ba, ATP synthase subunit beta, probable tubulin beta chain, prohibitin and RNA recognition protein were detected as DENV-2 binding proteins in brush border membrane fraction of *D. melanogaster* (Fig. 1; Table 2, Supplementary Information Table 2). The identification of protein bands were further confirmed using monoclonal antibodies. DENV-2 interacting proteins documented earlier and the observations of current study suggest that *Drosophila* possesses the necessary molecules which could help DENV-2 in establishing the infection (Supplementary Information Table S1 for DENV binding proteins in insect cells). Therefore, it would be useful to test the susceptibility of *D. melanogaster* to DENV-2 by injecting non lethal dose of DENV-2 in thoracic region of adult flies [Exposure of *D. melanogaster* flies to a low dose of DENV-2 (~1 μl of 8.23×10^6^ PFU/ml) did not have an effect on life span nor increased mortality compared to controls Logrank test *P*=0.73 data not shown].

**Fig. 1.**
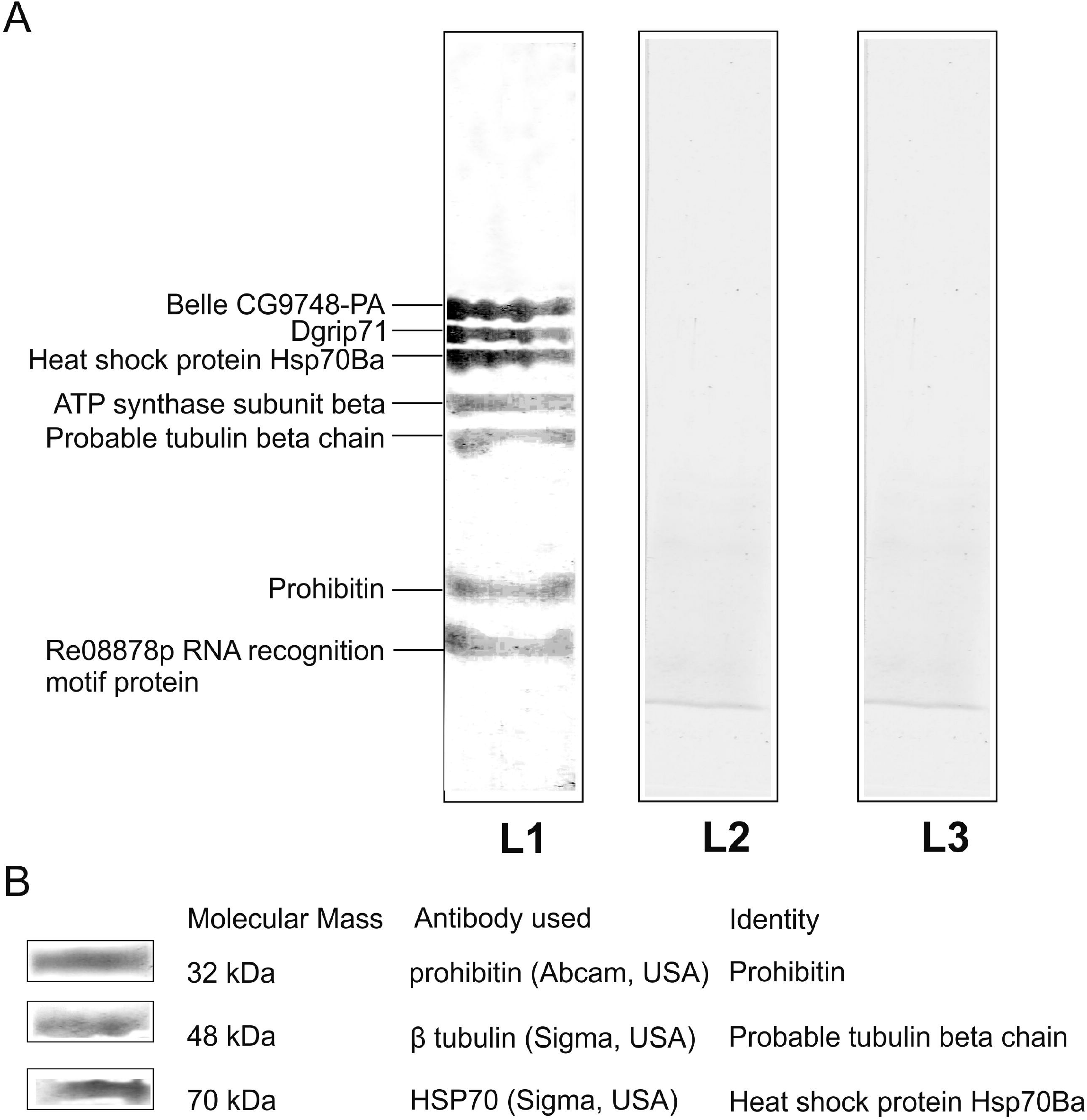
DENV-2 binding proteins in *D. melanogaster*. A) Membrane proteins from *D. melanogaster* midgut (lane L1, lane L2 and lane L3) were subjected to SDS–12.5% PAGE, transferred to nitrocellulose membrane, and incubated with DENV-2 (lanes L1, lane L2) and PBS (lane L3) at 37°C for 1 hour. The putative Dengue virus interacting proteins revealed after incubation with the anti-dengue-2 rabbit antibody (Lane L1 and Lane L3) and with a second antibody, an antibody mouse anti rabbit IgG conjugated to peroxidase. Lane B was only incubated with secondary antibody. Color was developed with H_2_O_2_ and DAB. B) Verification of MALDI TOF/TOF results using monoclonal antibodies Membrane proteins were subjected to 12.5% SDS PAGE and transferred to nylon membrane. Anti HSP70 antibody, Anti tubulin antibody and anti prohibitin antibody detected the bands corresponding to 70 kDa, 48 kDa and 32 kDa respectively. Anti actin antibody detected band corresponding to 42 kDa. However, 42 kDa band was not detected in VOPBA.

**Table 2:** Molecular identification of DENV-2 binding proteins from *D. melanogaster* midgut.

| No. | Accession<br>No. | Protein Description | Mol. Mass (kDa) |  | Mass<br>values<br>matched |
| --- | --- | --- | --- | --- | --- |
|  |  |  | From | From |  |
|  |  |  | Figure 1 | Database |  |
| 1 | **<u>NP_536783</u>** | Belle CG9748-PA | 84 | 85.029 | 10 |
| 2 | **<u>AAO49246</u>** | Gamma-tubulin ring complex<br>subunit Dgrip71 | 78 | 71.704 | 14 |
| 3 | **<u>AAK67155</u>** | Heat shock protein Hsp70Ba* | 70 | 70.191 | 10 |
| 4 | **<u>Q05825</u>** | ATP synthase subunit beta | 55 | 54.074 | 13 |
| 5 | **<u>Q9VRX3</u>** | Probable tubulin beta chain* | 48 | 50.697 | 7 |
| 6 | **<u>NP_724165</u>** | Lethal (2) 37Cc CG10691-PA,<br>isoform A (Prohibitin)* | 32 | 30.384 | 13 |
| 7 | **<u>AAN71293</u>** | RE08878p RNA recognition<br>motif protein | 26 | 28.007 | 7 |

\* The monoclonal anti-prohibitin antibody (Abcam, USA) interacted with 32-kDa protein, anti-tubulin antibody (Sigma Aldrich, Germany) recognized 48 kDa protein and anti-HSP70 antibody (Abcam, USA) recognized the 70-kDa protein in brush border membrane fractions of *D. melanogaster* midgut.

### Susceptibility of <u>D.</u> <u>melanogaster</u> to DENV-2

To determine the DENV-2 susceptibility, a non lethal dose of DENV-2 virus (~1 μl of 8.23×10^6^ PFU/ml) was injected in thoracic region of adult *Drosophila* flies. DENV-2 was detected in *D. melanogaster* midgut, brain and carcasses every day till 10 days p.i.. Immuno-fluorescence microscopy showed the presence of DENV-2 in *D. melanogaster* brain after 7 days of post infection (Fig. 2A). Detection of virus in gut tissue in addition to its time dependent appearance in brain indicated propagation of virus in the body of *D. melanogaster* flies (Fig. 2A). RT-PCR results showed the presence of DENV-2 in carcasses of *D. melanogaster* (Fig. 2B). Sequencing of RT-PCR product confirmed the specificity of RT-PCR. Susceptibility of *D. melanogaster* (61%±6.4) to DENV-2 at 7 days p.i. was found to be comparable to *Ae. aegypti* (72%±13) (Mann-Whitney U test *P*>0.05) (Fig. 2C).

**Fig. 2.**
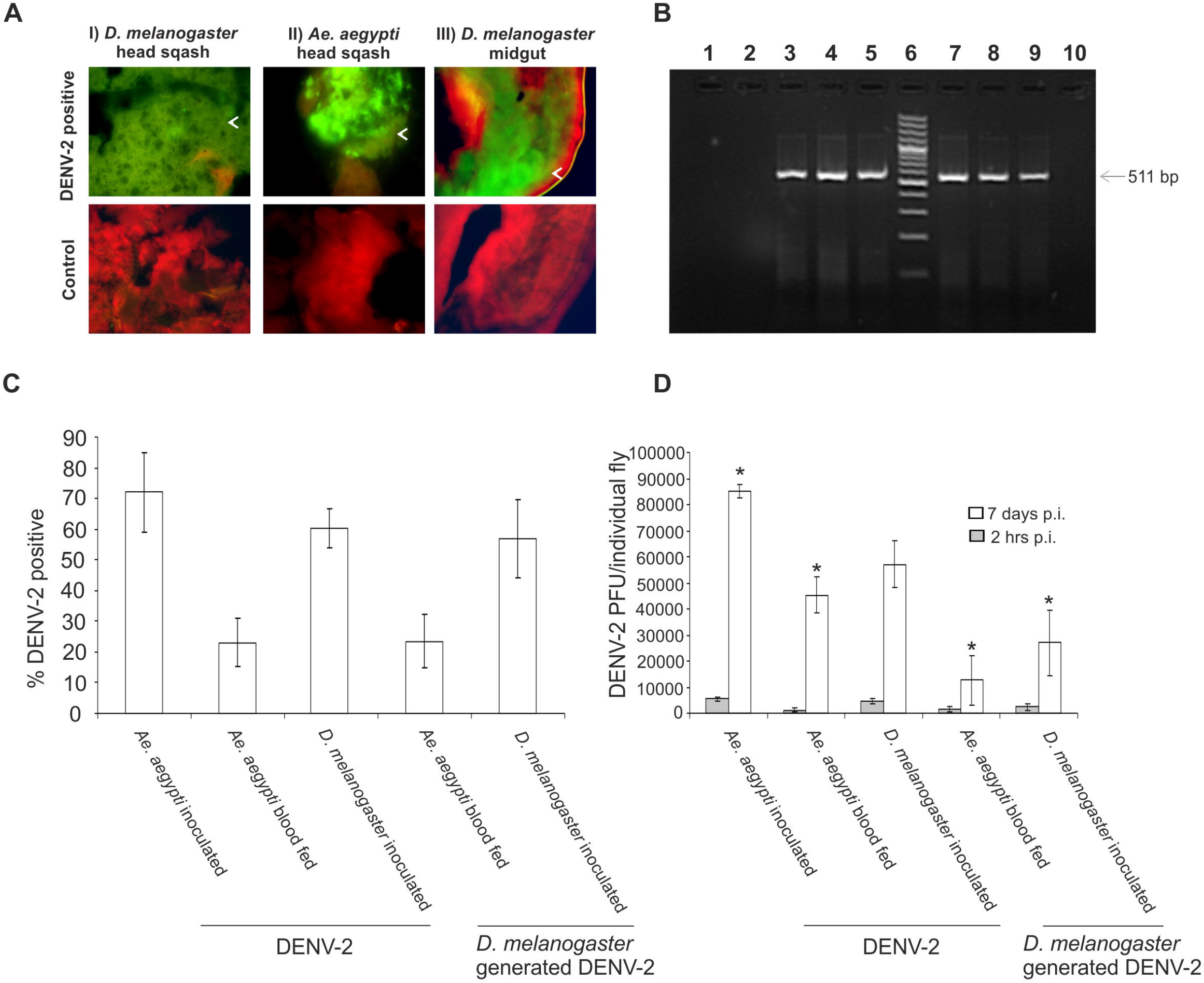
Susceptibility of *D. melanogaster* to DENV-2. A) Detection of DENV-2 in head squash of *D. melanogater* and *Ae. aegypti: D. melanogaster* and *Ae. aegypti* were intra thoracically injected with DENV-2 (3-4 μl of 10^6^ PFU /ml) or PBS (pH 7.4). DENV-2 was detected in head squash at 7 days p.i. using immuno-fluorescence microscopy. DENV-2 stained with FITC conjugated antibody (green color) and the neural tissue mass with Evan’s Blue (red color). I) *D. melanogaster* head squash, II) *A. aegypti* mosquitoes flies III) Midgut of *D. melanogaster*. B) RT-PCR detection of DENV-2: DENV-2 detected in carcasses of *D. melanogaster* and *Ae. aegypti* using RT-PCR. 1) Uninfected *Ae. aegypti* 2) Uninfected *D. melanogaster* 3) Oral infection DENV-2 in *Ae. aegypti* 4) intra thoracic injection of DENV-2 in *Ae. aegypti* 5) intra thoracic injection of DENV-2 in *D. melanogaster* 6) DNA ladder 100bp 7) intra thoracic injection of homogenate of DENV-2 positive *D. melanogaster* in *Ae. aegypti* 8) oral feeding of homogenate of DENV-2 positive *D. melanogaster* in *Ae. aegypti* 9) intra thoracic injection of homogenate of DENV-2 positive *D. melanogaster* in *D. melanogaster* 10) non template control (NTC). C) *Ae. aegypti* and *D. melanogaster* susceptibility to DENV-2: Percent DENV-2 positive head squash preparations of mosquitoes and flies were detected by immunofluorescence assay. DENV-2 inoculated *Ae. Aegypti* and *D. melanogaster* flies showed similar pattern of DENV-2 susceptibility (Mann-Whitney U test *P*>0.05) D) DENV-2 quantitation using plaque assay: DENV-2 titers in carcasses of individual *D. melanogaster* and *Ae. aegypti* at 2 hrs p.i. and 7 days p.i were quantitated using plaque assay.* significant difference in plaque forming units as compared to 2hrs p.i..

### Determination of infectivity of <u>D</u>. <u>melanogaster</u> generated DENV-2

In order to determine infectious nature of DENV-2 produced in *D. melanogaster*, both *D. melanogaster* and *Ae. aegypti* were infected with homogenates of DENV-2 virus positive flies. In orally fed *Ae. aegypti*, the DENV-2 antigen was detected in head squash preparations after 11 days (24%), while 56 % of inoculated *D. melanogaster* showed presence of DENV-2 in brain after 7 days (Fig. 2C). These data not only confirmed the *D. melanogaster* susceptibility to DENV-2, but also demonstrated that the infectious nature of *D. melanogaster* generated DENV-2. These experiments were repeated several times with reproducible results.

### DENV-2 quantitation using plaque assay

Plaque assays were used to determine the multiplication of DENV-2 in *Ae. Aegypti* and *D. melanogaster*. *Ae. aegypti* mosquitoes were infected with DENV-2 by oral feeding and intra-thoracic inoculation and *D. melanogaster* flies were infected intra thoracically with DENV-2 and were maintained at 28°C for 7 days. The DENV-2 titers in the carcasses of DENV-2 positive insects were determined by plaque assay (Fig. 2D). No significant difference was observed in DENV-2 viral load at 2 hrs p.i. in DENV-2 inoculated *Ae. aegypti* (5408±862 PFU/mosquito) and *D. melanogaster* (4511±968 PFU/fly) (Mann-Whitney U test *P*>0.05). These observations suggest that similar dose of DENV-2 was given to *Ae. aegypti* and *D. melanogaster*. At 7 days p.i., as compared to 2 h p.i., DENV-2 viral load was significantly increased in *Ae. aegypti* (85000±2684 PFU/mosquito) (Mann-Whitney U test *P*<0.05) and *D. melanogaster* (56983±8962 PFU/fly) (Mann-Whitney U test *P*<0.05). The homogenates of DENV-2 positive *D. melanogaster* was used to infect *D. melanogaster* and *Ae. aegypti*. At 7 days p.i., as compared to 2 h p.i., the DENV-2 viral load was significantly increased in oral fed *Ae. aegypti* (12569±9638 PFU/mosquito) (Mann-Whitney U test *P*<0.05) and intra thoracic injected *D. melanogaster* (26982±12692 PFU/fly) (Mann-Whitney U test *P*<0.05). At 7 days p.i., the virus titer in intra thoracic injected *D. melanogaster* and *Ae. Aegypti* flies was almost similar (Mann-Whitney U test *P*>0.05).

### Anti-DENV-2 response in <u>D</u>. <u>melanogaster</u>

The antiviral response of *D. melanogaster* against the DENV-2 virus infection was checked in selected antiviral pathway genes using the qPCR assays. The expression levels of key components of JAK-STAT, RNA interference (RNAi) and Toll pathway were assessed in DENV-2 infected *D. melanogaster* using qPCR at 24h, 48h and 7 days p.i.. DENV-2 stimulates the transcriptional activation of RNAi, JAK-STAT and Toll pathway (Table 3). Transcript levels of key mediators of RNAi pathway, Argonaute 1, Argonaute 2, Argonaute 3, Dicer 1, Dicer 2, Drosha and Pasha were up-regulated during course of DENV-2 infection (Mann-Whitney U test *P*<0.05). ADAR, FMR, Loqs, RNBP and VIG showed slight variation in transcript levels in response to DENV-2 infection (Table 3). Transcript levels of JAK, STAT, Prohitibin, Rel1 and Toll were up-regduring DENV-2 infection (Table 3) (Mann-Whitney U test *P*<0.05).

**Table 3:**
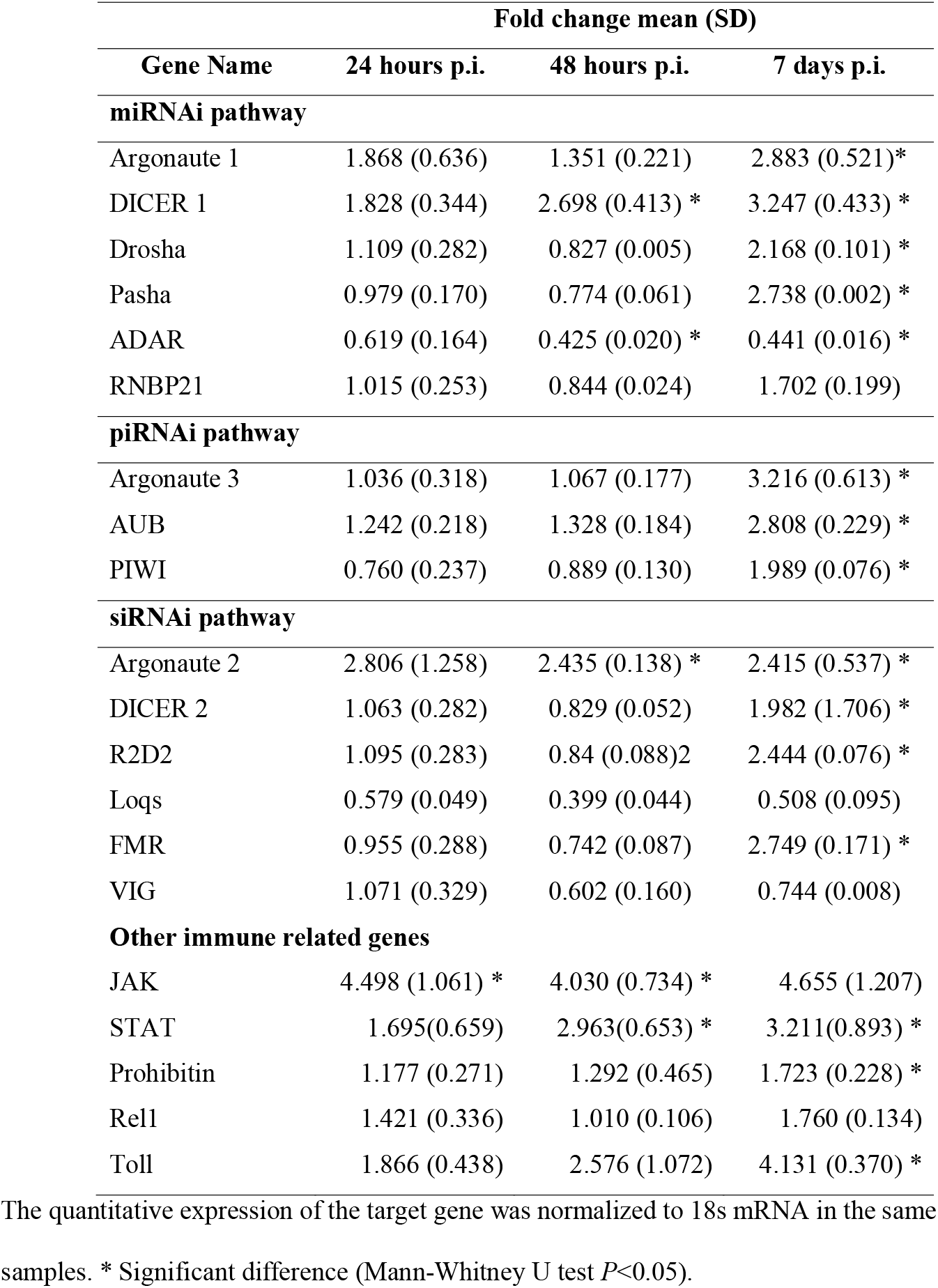
Relative gene expression changes in *D. melanogaster* in response to DENV-2 infection.

| Gene Name | Fold change mean (SD) |  |  |
| --- | --- | --- | --- |
|  | 24 hours p.i. | 48 hours p.i. | 7 days p.i. |
| **miRNAi pathway** |  |  |  |
| Argonaute 1 | 1.868 (0.636) | 1.351 (0.221) | 2.883 (0.521)* |
| DICER 1 | 1.828 (0.344) | 2.698 (0.413) * | 3.247 (0.433) * |
| Drosha | 1.109 (0.282) | 0.827 (0.005) | 2.168 (0.101) * |
| Pasha | 0.979 (0.170) | 0.774 (0.061) | 2.738 (0.002) * |
| ADAR | 0.619 (0.164) | 0.425 (0.020) * | 0.441 (0.016) * |
| RNBP21 | 1.015 (0.253) | 0.844 (0.024) | 1.702 (0.199) |
| **piRNAi pathway** |  |  |  |
| Argonaute 3 | 1.036 (0.318) | 1.067 (0.177) | 3.216 (0.613) * |
| AUB | 1.242 (0.218) | 1.328 (0.184) | 2.808 (0.229) * |
| PIWI | 0.760 (0.237) | 0.889 (0.130) | 1.989 (0.076) * |
| **siRNAi pathway** |  |  |  |
| Argonaute 2 | 2.806 (1.258) | 2.435 (0.138) * | 2.415 (0.537) * |
| DICER 2 | 1.063 (0.282) | 0.829 (0.052) | 1.982 (1.706) * |
| R2D2 | 1.095 (0.283) | 0.84 (0.088)2 | 2.444 (0.076) * |
| Loqs | 0.579 (0.049) | 0.399 (0.044) | 0.508 (0.095) |
| FMR | 0.955 (0.288) | 0.742 (0.087) | 2.749 (0.171) * |
| VIG | 1.071 (0.329) | 0.602 (0.160) | 0.744 (0.008) |
| **Other immune related genes** |  |  |  |
| JAK | 4.498 (1.061) * | 4.030 (0.734) * | 4.655 (1.207) |
| STAT | 1.695(0.659) | 2.963(0.653) * | 3.211(0.893) * |
| Prohibitin | 1.177 (0.271) | 1.292 (0.465) | 1.723 (0.228) * |
| Rel1 | 1.421 (0.336) | 1.010 (0.106) | 1.760 (0.134) |
| Toll | 1.866 (0.438) | 2.576 (1.072) | 4.131 (0.370) * |

The quantitative expression of the target gene was normalized to 18s mRNA in the same samples. \* Significant difference (Mann-Whitney U test $P < 0.05$ ).

## DISCUSSION

Availability of large volume of information on genetics, development and genome of *D. melanogaster* as well as its potential to use as a test system to analyze human diseases^12–18^, makes this non vector dipteran as an attractive model to study the pathogens and propagation in these pathogens in the insect model systems ^19–24,27^. Using genome-wide RNA interference screen in *D. melanogaster* the insect host factors required for DENV propagation were identified^25^. The results of Sessons et al. study indicate that remarkable conservation in required factors between the dipteran and human hosts. Recently it has been demonstrated that RNA interference modulates the replication of dengue virus in *D. melanogaster* cells^25,32^. Results obtained in these studies clearly suggest that DENV could propagate in Drosophila cells. Recently Rences et al.^33,34^ utilized the *D. melanogaster* and *Ae. aegypti* to understand the molecular mechanisms involved in *Wolbachia-* mediated pathogen protection. The infection with *Wolbachia* efficiently reduced the DENV replication in *D. melanogaster* as well as in *Ae. aegypti* ^33,34^. The results obtained in these studies demonstrated that the mechanism of DENV blocking by *Wolbachia* is more complicated than a simple priming of the insect innate immune system. It will be interesting to investigate the complex mechanism involved in DENV-vector interactions. Availability of *Drosophila* mutants will help in deciphering the complex interactions involved in DENV infection. The binding proteins, susceptibility and virus titer of DENV-2 in *D. melanogaster* was not investigated in earlier studies. We, therefore, infected *D. melanogaster* with DENV-2 and found that *D. melanogaster* was not only susceptible to DENV-2 but also produced infectious DENV-2 particles. The DENV-2 infection in *D. melanogaster* seems to be a specific pathogenic process rather than nonspecific viremia. First, DENV-2 was detected in midgut, carcasses and brain after 7 days of post infection period in *D. melanogaster* similar to in *Ae. aegypti*. Second, DENV-2 virus produced in fly is equally infectious to both *D. melanogaster* and *Ae. aegypti*. Third, after 7 days post infection DENV-2 titers were comparable in *Ae. aegypti* and *D. melanogaster*. These qualities make *D. melanogaster* a potential model system for examining DENV-vector interactions. The Drosophila model system is useful in rapid and unbiased identification of host factors involved in pathogenesis^12,14,19,26,27^. Considering the biosafety issues, the fruit fly system has certain added advantages in comparison with the anthropogenic vectors such as mosquito, sand fly and tick.

DENV-2 binding proteins in mosquito cells have been identified in several studies^2,3,5,6^ [see supplementary Information Table 1]. However, the identity of most of the proteins is not conclusively determined. Proteins such as HSP 70Ba, ATP synthase sub unit beta, probable tubulin beta chain and prohibitin were found interacting with *Ae. aegypti* and *D. melanogaster*. Based on results obtained in the previous studies and current study, it is reasonable to infer that DENV-2 may share the similar receptor molecule(s) on dipteran cells^2,3,5^. HSP 70 proteins might be binding receptor and other proteins such as actin, tubulin, ATP synthase etc. might be secondary receptors. Although in most cases, individual viruses have their own distinct and specific receptors, in some cases the same set of receptors can be used by many different viruses. Japanese Encephalitis Virus (JEV), DENV and West Nile Virus (WNV) share the similar molecules for their entry into the mosquito cells^2,3,5,6,35,36^. Perhaps the best studied example of this is the HSP 70 of *Ae. aegypti*, which is used by DENV, JEV and WNV as a receptor^2,5,6,7^. As these molecules are house keeping in nature, they are present in other dipteran insects also. We hypothesize that DENV-2 might share similar receptors in dipteran insects such as *Ae. aegypti, Ae. albopictus* and *D. melanogaster*.

Interestingly in VOPBA experiments, Belle and RNA recognition protein were recognized as DENV-2 binding proteins. In *Drosophila*, Belle, a DEAD-box RNA helicase, has been documented to regulate RNA interference (RNAi)^37^. During the infection, interaction and co-localization of DDX3 (a human homologue of Belle) with arboviral proteins and viral RNA have been demonstrated ^38,39^. Similarly, the role of RRM proteins in the post-transcriptional gene expression modulation of the *Drosophila* RNAi pathway is well documented^37^. These observations hint the possible involvement of *Drosophila* RNAi pathway in controlling the DENV-2 infection. Prohibitins (PHBs) are highly conserved proteins in eukaryotes and are associated with various cellular functions including the immune regulation. Role of prohibitin as a non-receptor interacting polypeptide in DENV-2 infection has been reported previously^2,29^. These observations suggest that similar immune response might be triggered in *Ae. aegypti* and *D. melanogaster* to counteract the DENV-2 infection.

In the mosquito, arboviruses are confronted with RNAi, JAK-STAT and Toll pathway ^10–12,40–49^. RNAi is one of the important innate antiviral pathways in *Ae. aegypti* that controls the arbovirus replication^41,42,44–47,49–58^. The recent reports demonstrated that RNAi pathway can eliminate the DENV from *Ae. aegypti* and Dcr-2 and Ago-2 knockdowns enhances the DENV replication^42,45,47,49^. Similarly it has been also demonstrated that DENV can grow in *Drosophila* S2 cells and the RNAi regulates the DENV replication in these cells^32^. Therefore, we employed the *Drosophila* RNAi, JAK-STAT and Toll gene expression to address how a model innate system responds to DENV-2 infection. Transcript levels of most of the key mediator of RNAi response were up-regulated in response to DENV-2 infection at 7 days p.i.. Moderate modulation in RNAi pathway genes was observed at early time points 24 hrs and 48 hrs p.i.. Loqs expression was slightly down-regulated in response to DENV-2 infection. It has been demonstrated that dsRNA of viruses can be cleaved by Dcr-2 without Loqs-PD and complete knock down of Loqs-PD has no effect on antiviral silencing^37,59^. These observations suggest that Loqs have limited role in antiviral response in *Drosophila*.

Recent transcriptome analysis of DENV2-infected *Ae. aegypti* reveled the involvement miRNA pathway in virus infection^51–53^. The significant increase in unique miRNAs was observed during DENV infection in *Ae. aegypti*^58^. Over the course of infection, 9 days p.i. time point showed maximum number of unique modulated miRNAs^58^. It has been suggested at 9 days p.i. the repair mechanisms in uninfected mosquitoes is activated and results in to significant increase in the miRNA levels at time point^58^. The expression of Argonate-1, Dicer-1, Drosha and Pasha increased during the time course of DENV infection in *D. melanogaster*. These observations suggest that miRNA pathway activity is altered during DENV-2 infection in *D. melanogaster*. The results obtained in current study corroborate with earlier studies that showed the presence of modulated miRNAs in DENV2 infection in mosquitoes.

A significant role of piRNA pathway has been also reported in various arbovirus infections^54,55,57^. It has been envisaged that a non-canonical piRNA pathway play important role in vector mosquitoes and target alphavirus replication^54^. The activity PIWI proteins and virus-specific piRNA molecules is also detected in somatic cells of *D. melanogaster* suggesting that the piRNA pathway plays important role in antiviral functions in insects^50^. Induction in transcript levels of Argonaute 3 AUB and PIWI suggest that DENV-2 replication is able to trigger the piRNA pathway. The miRNA, piRNA and siRNA pathways may act in amalgamation to control DENV-2 infection in *D. melanogaster*.

The up-regulation of JAK, STAT, REL1 and Toll proteins suggested the activation of the Toll pathway and JAK-STAT pathway (Table 3). In *Ae. aegypti*, the JAK-STAT and Toll pathway are involved in the anti-dengue defense^10^. Up-regulation of the Toll pathway has also been reported in *Ae. aegypti* in response to Sindbis virus infection^40^. The fruit fly seems to rely on RNAi, JAK-STAT and Toll to counteract DENV-2 infections. Though the JAK-STAT, RNAi and Toll pathways were seen to be induced in response to DENV infection, 60% of flies were still infected by DENV-2. These observations suggest that JAK-STAT, RNAi and Toll pathway are activated in response to DENV-2 infection but are not sufficient for complete elimination of DENV-2 in *Drosophila*. DENV-2 must have evolved strategies to counteract the effects of the JAK-STAT, RNAi and Toll pathway.

In depth studies on host factors involved in virus infection is necessary to design and develop effective intervention strategies. In many other insect vectors such as ticks, sand flies, fleas and other members of Culicidae family, amenable genetic systems limit in depth understanding of vector parasite interaction in these species. In this context, *Drosophila* becomes an attractive model system to elucidate the complex host-parasite interactions. *Drosophila* possesses the necessary repertoire of proteins that might require for virus entry. *D. melanogaster* supports the growth of WNV, SINV and DENV-2. Further the immune response mounted against arboviruses is similar in *D. melanogaster* and *Ae. aegypti*. Due to whole genome sequence and established techniques for genetic and developmental manipulations, *D. melanogaster* turn out to be an attractive model organism to understand molecular and cellular mechanisms in host-arbovirus interactions. *D. melanogaster* could be used as surrogate invertebrate host model system and can be used to study parasite-vector interactions in less characterized vectors such as ticks, sand flies, fleas and other Culicine mosquitoes.

## CONCLUSION

In conclusion, VOPBA revealed that DENV-2 exploit similar molecules in *D. melanogaster* and *Ae. aegypti* for its entry. *D. melanogaster* supports the growth of DENV-2 virus and *D. melanogaster* generated DENV-2 was able to infect the *Ae. aegypti* with similar kinetics. Results obtained in this study and earlier reports suggest that *D. melanogaster* and *Ae. aegypti* mount similar immune response against the invading arboviruses. These qualities make *D. melanogaster*, a potential model system for the study of arbovirus-vector interactions.

## ACKNOWLEDGEMENTS

We thank the Director, National Institute of Virology, Pune for the providing the facilities. We thank Dr. Dipankar Chaterjee, Indian Institute of Science, Bangalore for MALDI TOF/TOF analysis. This research was supported by UGC-CAS to Department of Zoology, University of Pune and ICMR grant to Prof. Dileep Deobagkar.

**Supplementary Information Table S1**: DENV-2 interacting proteins in insect cells

**Supplementary Information Table S2**: Molecular identification of DENV-2 binding proteins from *D. melanogaster* midgut using 1D and 2D VOPBA

