## Supplementary Table S1 for "*Drosophila melanogaster* as a model host to study arbovirus–vector interaction"

**Supplementary information Table S1: Dengue binding proteins in insect cells**

| **Cell line/species** | **Cell / tissue type** | **Dengue Virus Serotype** | **Receptor Characteristics** | **Type of Assay** | **Reference** |
| --- | --- | --- | --- | --- | --- |
| C6/36 | *Ae. albopictus* cell line | Dengue-4 stain H-241 | Two glycoproteins 40/45 kDa | VOPBA | Salas-Benito and del Angel 1997 |
| *Ae. Aegypti* organs | Midgut, salivary glands from adults as well as egg, larvae and pupa cell extract | Dengue-4 stain H-241 | A 45 kDa glycoprotein | VOPBA | Salas-Benito and del Angel 1997 |
| C6/36 | *Ae. albopictus* cell line | Dengue-2 New Guinea C strain | 80 and 67 kDa protein | VOPBA | Munoz et al 1998 |
| C6/36 | *Ae. albopictus* cell line | Dengue-4 | 45 kDa protein | VOPBA | Yazi-Mendoza et al., 2002 |
| C6/36 | *Ae. albopictus* cell line | Dengue-2 New Guinea C strain | Tubulin β chain protein | VOPBA | Chee and AbuBakar 2004 |
| C6/36, *Ae. Aegypti* | *Ae. albopictus* cell line, *Aedes aegypti* midgut | Dengue-1, strain Hawaii; Dengue-2, strain New Guinea C (NGC); Dengue-3, strain H-87; and Dengue-4, strain H-341. | R80, R67, R57 | VOPBA | Mercado-Curiel et al., 2006 |
| C6/36 | *Ae. albopictus* cell line | Dengue-2 | 74 kDa and 45 kDa glycoprotein | Affinity Chromatography | Salas-Benito et al., 2007 |
| *Ae. Aegypti DS3*, *DMEB* and *IBO-11* strains | *Ae. aegypti* midgut | Dengue-2 Jamaica strain | R67/R64 | VOPBA | Mercado-Curiel et al., 2008 |
| *Ae. aegypti* | *Ae. aegypti* salivary gland extract | Dengue 1-4 reference strain | 77, 58, 54 and 37 kDa protein | VOPBA | Cao-Lormeau 2009 |
| *Ae polynesiensis* | *Ae polynesiensis* salivary gland extract | Dengue-1 and Dengue-4 | 67, 56, 54, 50 and 48 kDa | VOPBA | Cao-Lormeau 2009 |
| C6/36, A7 cells *Ae. aegypti* | *Ae. aegypti midgut, Ae. albopictus* cell line, *Ae. aegypti* cell line | Dengue-2 | Actin, ATP synthase β subunit, HSc 70, orisis, prohibitin, tubulin β chain, and vav-1 | VOPBA | Paingankar et al., 2010 |
| C6/36, CCL-125 and adult *Ae. aegypti* mosquito cells, | *Ae. aegypti, Ae. albopictus* cell line | Dengue-2 | Prohibitin | VOPBA | Kuadkitkan et al., 2010 |
| C6/36 | *Ae. albopictus* cell line | Dengue-4 | 70 kDa Heat Shock and 70 kDa Heat Shock cognate proteins (HSP70/HSc70), Binding immunoglobulin protein (BiP), Thioredoxin/protein disulphide isomerase (PDI), and 44 kDa Endoplasmic reticulum resident protein (ERp44) | Affinity Chromatography | Vega-Almeida et al., 2013 |
